# Cortical layer-specific modulation of neuronal activity after sensory deprivation due to spinal cord injury

**DOI:** 10.1101/2020.12.28.424612

**Authors:** Marta Zaforas, Juliana M. Rosa, Elena Alonso-Calviño, Elena Fernández-López, Claudia Miguel-Quesada, Antonio Oliviero, Juan Aguilar

**Author notes:** These authors contributed equally (Marta Zaforas, Juliana M. Rosa). This article was first published as a preprint: Zaforas M*, Rosa JM*, Alonso-Calviño E, Fernández-López E, Miguel-Quesada C, Oliviero A, Aguilar J (2021). Cortical layer-specific modulation of neuronal activity after sensory deprivation due to spinal cord injury. bioRxiv. https://doi:10.1101/2020.12.28.424612.

## Abstract

Cortical areas have the capacity of large-scale reorganization following sensory deprivation. However, it remains unclear whether this phenomenon is a unique process that homogenously affects an entire deprived cortical region or it is suitable to changes depending on neuronal networks across distinct cortical layers. Here, we studied how local circuitries within each layer of the deprived cortex set the basis for neuroplastic changes after immediate sensory deprivation due to thoracic spinal cord injury (SCI) in anaesthetised rats. *In vivo* electrophysiological recordings from deprived hindlimb somatosensory cortex showed that SCI induces layer-specific changes mediating evoked and spontaneous activity. In supragranular layers 2/3, sensory deprivation increased gamma oscillations and the ability of these neurons to initiate up-states during spontaneous activity, suggesting altered corticocortical network and/or intrinsic properties that may serve to maintain the excitability of the cortical column after deprivation. On the other hand, sensory deprivation enhanced infragranular layers’ ability to integrate evoked-sensory inputs leading to increased and faster neuronal responses. Delayed evoked-responses onset were also observed in layers 5/6, suggesting alterations in thalamocortical connectivity. Altogether, our data indicate that SCI immediately modifies local circuitries within the deprived cortex allowing supragranular layers to better integrate spontaneous corticocortical information, and thus modifying column excitability, and infragranular layers to better integrate evoked-sensory inputs to preserve subcortical outputs. These layer-specific neuronal changes may guide the long-term alterations in neuronal excitability and plasticity associated to the rearrangements of somatosensory networks and the appearance of central sensory pathologies usually associated with spinal cord injury.

## INTRODUCTION

Spinal cord injury (SCI) produces an abrupt and robust loss of sensory inputs onto cortical areas that receive information from body regions below the lesion level (i.e. hindlimb cortex that receives afferent inputs from hindlimbs). This sensory deprivation initiates the so-called process of cortical reorganization (CoRe), which is typically described as an expansion of the neuronal activity from intact cortical areas towards the sensory deprived cortex (Curt et al., 2002; Endo et al., 2007; Ghosh et al., 2010; Jain et al., 1998, 2008). CoRe plays key roles in the functional recovery after SCI, but it has also been implicated in the generation of associated pathologies such as neuropathic pain and spasticity (Siddall et al., 2003; Wrigley et al., 2009). Similar cortical rearrangements after sensory deprivation have also been described in different sensory systems as visual (Griffen et al., 2017) and auditory cortex (Bola et al., 2017). Most of studies about cortical reorganization after SCI have used large-scale experimental approaches such as extracranial electroencephalographic recordings (Green et al., 1998), voltage sensitive dye (Ghosh et al., 2010) and functional magnetic resonance imaging (Endo et al., 2007) in a period ranging from days-to-months after the injury. However, it is currently known that such functional changes are observed immediately after SCI in different animal models as rodents (Aguilar et al., 2010; Humanes-Valera et al., 2013; Yagüe et al., 2014; Yague et al., 2011) and pigs (Jutzeler et al., 2019). In this line, our group have previously shown that both intact and deprived somatosensory cortex became more responsive to peripheral sensory stimulation above lesion level few minutes after the lesion (Humanes-Valera et al., 2013; Yagüe et al., 2014), while spontaneous activity is drastically reduced (Aguilar et al., 2010; Fernández-López et al., 2019). In addition to the cortical changes, SCI also modifies thalamic and brainstem spontaneous and evoked neuronal excitability, which may be linked to the changes observed in the somatosensory cortex (Alonso-Calviño et al., 2016; et al., Jain et al., 2008).

Changes in the cortical activity after SCI have been mostly studied by using electrophysiological recordings from layer 5 neurons. However, the neocortex is a complex structure composed of six layers organized in distinct vertical columns. Within each layer, different functional properties and input/output connections are achieved by interconnected excitatory pyramidal neurons, inhibitory neurons, and glial cells. Although several studies in the last decade have shed light into distinct strategies used by each layer to encode information, less is known about the functional changes induced within each cortical layer following sensory deprivation that may precede longterm cortical reorganization and could help to understand the physiological changes in brain areas controlling the integration and processing of sensory inputs.

Using *in vivo* electrophysiological recordings from anesthetised rats, we studied how neuronal activity mediated by corticocortical and thalamic connections as well as local circuitries in the hindlimb cortex (HLCx) are immediately affected by sensory deprivation after a thoracic SCI. For that, we used a vertical multielectrode array to determine the neuronal excitability across layers of the deprived HLCx during evoked and spontaneous activity. Peripheral stimulation of the contralateral forelimb showed a layer-dependent increase in sensory evoked-LFP responses across HLCx layers indicating that changes in neuronal network properties of the deprived cortical column may favour the excitability. However, a striking heterogeneity was observed when other physiological parameters were analysed. Infragranular L5/6, but not L2/3, exhibited increased LFP slope, increased multiunit activity and delayed onset latencies. On contrary to evoked-responses, spontaneous activity was mostly affected in supragranular layers as observed by increased high-rhythms frequencies and probability to initiate UP-states. Altogether, our data indicate that SCI immediately modifies local circuitries within the deprived cortex allowing supragranular layers to better integrate spontaneous corticocortical information, thus modifying the excitability of the column, and infragranular layers to better integrate evoked-sensory inputs to preserve subcortical outputs.

## METHODS

### Ethical Approval

Experiments were performed on male Wistar rats (n = 36, Ctr:WI Charles-River RRID:RGD_737929) aging 2-6 months, mean weight 395 g. (SD 45). Animals were housed 2 per cages in standardized cages, with *ad libitum* access to food and water and maintained at 23 °C on a 12-hour light/dark cycle. All experiments were performed in accordance with the International Council for Laboratory Animal Science and the European Union 2010/63/EU, ARRIVE guidelines and comply with the policies and regulations set out by *The Journal of Physiology*. All steps of the experimental procedure were performed in a way to minimise the animals’ pain and suffering and were approved by the Ethical Committee for Animal Research at the Hospital Nacional de Parapléjicos (152CEEA/2016). After experimental procedure, animals were sacrificed by anaesthetic overdose (urethane 1.5 g/kg i.p., Sigma-Aldrich, U2500-100G). Approximately 14% of the total number of animals died during the experimental procedure of the spinal cord injury or before it, due to anaesthesia intolerance. All researchers involved in the study were aware of the ethical principles under which the Journal operates and complied with the animals’ ethics checklist set out by *The Journal of Physiology*.

### Experimental approach

The general experimental approach (anaesthesia, surgery and peripheral stimulation) was similar to that used in our previous studies (Aguilar et al., 2010; Alonso-Calviño et al., 2016; Humanes-Valera et al., 2013; Humanes-Valera et al., 2017). Briefly, animals were anaesthetised with an intraperitoneal injection of urethane (1.5 g/kg i.p.), placed in a stereotaxic frame (SR-6 Narishige Scientific Instruments, Tokyo, Japan) passively ventilated at 2 l O_2_/min by a mask (Medical Supplies & Services, INT. LTD., England) and body temperature kept constant at 36.5°C using a homeothermic blanket (Cibertec SL, Madrid, Spain). The optimal level of anaesthesia was constantly proved by lack of motor reflex after tail and limb pinch. Then, lidocaine 5% (Normon, Cat#P06B1) was applied subcutaneously into the areas of the incision and thoracic laminectomy (at T9–T10 vertebra) was performed keeping the dura mater intact and protected until the moment of performing a complete transection of the spinal cord. Next, the skull was exposed, and a craniotomy was performed on the right hemisphere over the hindlimb representation of the primary somatosensory cortex (AP 0 to −3 mm; ML 1 to 4 mm; Paxinos and Watson, 2007) to allow lowering a vertical array for further record of neuronal activity. The stability of recordings was improved by drainage of the cisterna magna and covering the exposed cortex with agar at 4% (Sigma-Aldrich, Cat#A7002-250G). The exact location of the probe was optimized by assessing the responses to tactile stimulation of the rat’s hindlimb with a cotton swab while listening to the recorded signal through a pair of loudspeakers.

### Electrophysiological recordings and peripheral stimulation

Extracellular recordings were obtained from 24 rats by a linear vertical probe of 32 iridium contacts with diameter 177 μm^2^ spaced at 50 μm (impedance 1-4 MΩ at 1 kHz; ref: A1×32-6mm-50-177-A32, NeuroNexus Technologies Inc., US). The array was slowly introduced (1-2 μm/s) through the craniotomy into the HLCx (Fiáth et al., 2019) and a ground electrode was placed in the parietal muscular tissue. The reference electrode was built in the vertical probe, 0.5 mm above the superficial recording site and outside the cortex (diameter 4200 μm^2^). Recording protocol started ~40 min after the end of the electrode insertion to allow recovery of cortical tissue following time line in Figure 1A. Spontaneous activity was recorded during 10 min. Stimulation protocol (0.5 ms duration at 0.5 Hz) was applied through bipolar needle electrodes (30G, B. Braun Melsungen AG.) located subcutaneously in the wrist of contralateral forelimb and hindlimb extremities. Two different intensities were applied: 1) low intensity (0.5 mA) to activate only a fraction of the available peripheral fibres, mainly low-threshold primary fibres running through the lemniscal pathway from the dorsal column to the brainstem, and 2) high intensity (5 mA) to activate the maximum number of fibres including high-threshold primary fibres that synapse in the dorsal horns of the spinal cord including the spinothalamic tract (Lilja et al., 2006; Yague et al., 2011). However, note that all data analysed throughout this study were obtained from responses showing an initial latency below 15 msec, which correspond to low-threshold peripheral fibers (tactile and proprioceptive) from the entire paw (digits and palm). After recordings of evoked and spontaneous activity in control conditions, complete transection of the spinal cord was performed using a spring scissors. Immediately after transection, pulses of 10 mA electrical stimulation were applied to the contralateral hindlimb to confirm that no physiological responses were evoked by stimuli delivered below the level of the lesion. Complete spinal cord transection was also visually confirmed under the surgical microscope by the total separation of the borders. Recordings were continuously acquired during the transection to confirm the stability of the recordings. Approximately 20-30 min after the transection, the same protocol as before SCI was applied. Based on the absence of reflexes to forelimb stimuli, spontaneous whisker movements and corneal reflex, animals never received additional anaesthesia between the pre lesion protocol and the post lesion protocol. All recording data were converted into digital data at a 40 kHz sampling rate (16/24 rats) and 1kHz (8/24 rats), with 16–bit quantization by an OmniPlex System controlled by OmniPlex Software (RRID:SCR_014803, Plexon Inc, Texas, USA). All the 40 kHz signals were offline filtered into two signals: local field potentials (LFP, low-frequency band: up to 1kHz) and multiunit activity (MUA, high-frequency band: 0.3-3kHz) by using Spike2.v7 (RRID:SCR_000903), Cambridge Electronics Design, Cambridge, UK.

**Figure 1:**
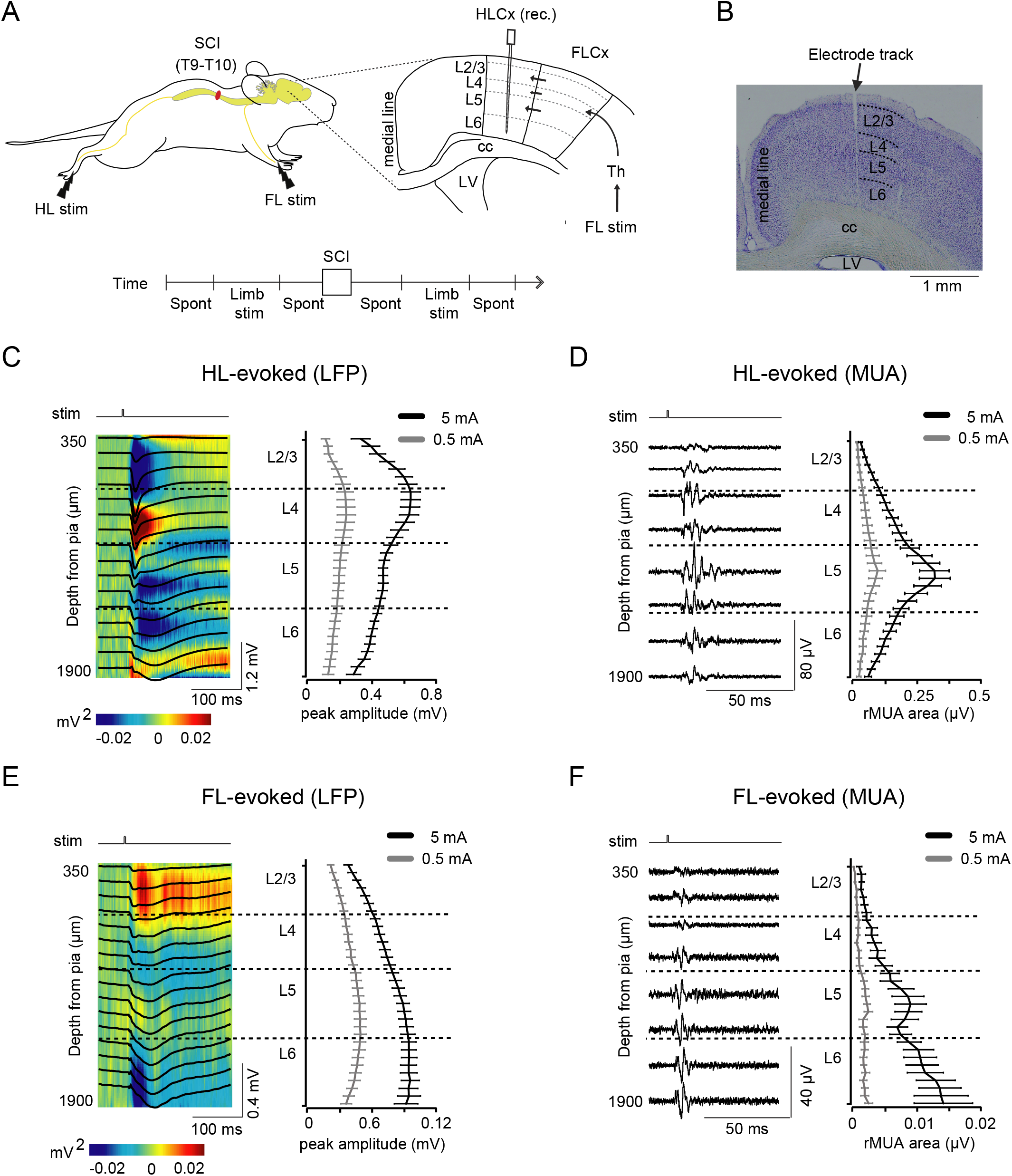
Experimental approach and laminar characterization of evoked sensory responses in hindlimb cortex. (A) Schematic illustration of the experimental protocol. Extracellular recordings were obtained using a multielectrode probe inserted in the hindlimb representation of the primary somatosensory cortex (HLCx) from anaesthetized rats. Complete transection of the spinal cord was performed at thoracic level (SCI, T9-T10). Spontaneous activity (Spont) and evoked responses to electrical stimulation at high (5 mA) and low (0.5 mA) intensity delivered to the contralateral hindlimb (HL stim) and forelimb (FL stim) were recorded following the described timeline in control conditions and immediately after SCI. In the cortical representation, black arrows indicate corticocortical connections and thalamic inputs into FL cortex in response to forepaw stimulation (FL stim). Cortical layers were designated as L2/3, L4, L5 and L6. Dashed black lines indicate boundaries of cortical layers. Corpus callosum, cc; Lateral ventricle, LV; Thalamus, Th. (B) Nissl-stained coronal section of a representative rat showing an electrode track in HL cortex. (C) Left: representative sensory evoked-LFP across HL cortical layers in response to a 5 mA electrical stimulation of the contralateral hindlimb (HL). Recordings obtained on every 100 μ m are shown on top of the current source density (CSD) map from the same recordings. Right: averaged evoked-LFP amplitudes as a function of cortical depth in response to low (0.5 mA, grey) and high (5 mA, black) sensory stimulation. (D) Left: High-filtered LFP traces from the same recordings in C showing evoked-MUA signal. Right: Averaged area of rMUA from the same population in C. (E) Left: Representative traces of evoked-LFP overlapped on the CSD map in response to 5 mA forelimb stimulation from the same animal as C-D. Note that in this case current sinks are stronger in L2/3 and L5. Right: averaged evoked-LFP amplitudes as a function of cortical depth in response to low (0.5 mA, grey) and high (5 mA, black) sensory stimulation. (F) Left: High-filtered LFP traces from the same recordings in E showing evoked-MUA signal. Right: Averaged area of rMUA from the same population in E. Dashed black lines indicate borders between layers. Errors bars represent SEM only for illustrative purposes (n = 24 rats). Population average and SD are given in Figure 3. See also - Supplemental Table 1.

### Data analysis: evoked responses and spontaneous activity

For laminar profile analysis, LFP evoked responses from each electrode were averaged across 100 stimuli (0.5 Hz) and measured as the maximum amplitude to negative peak (mV) in the local fast response in a time window corresponding to 5-60 ms or 5-30 ms following sensory stimulation of hindimb or forelimb, respectively. In order to quantify MUA, the filtered recordings were rectified (rMUA) and averaged across 100 stimuli to measure the total voltage resulting from the averaged area of responses (μV). The obtained value was then subtracted from the background voltage obtained 50 ms before stimulation. For layer analysis, LFP from electrodes within the same layer were obtained according to the depth: layer 2/3 (150-650 μm), layer 4 (700-1000 μm), layer 5 (1050-1450 μm) and layer 6 (1500-2000 μm; Fiáth et al., 2016). In the case of rMUA, neuronal signals obtained from individual channels within a layer were summed up to allow robust detection of the neuronal activity.

Onset latency of evoked-LFP and -rMUA was calculated for each layer by fitting the averaged response with an equation of the form of the Boltzmann charge–voltage function. This equation was solved for its 4th derivative giving a highly accurate measure of the response onset independent of the slope rise phase (Fedchyshyn and Wang 2007). Slopes were measured by using the next equation: 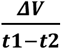, where ***ΔV*** is the LFP amplitude, t_1_ is the onset and t_2_ is the time of the negative peak. For analysis of the spontaneous cortical activity, up-states within the slow-wave activity (SWA) were first analysed in periods of 5 minutes of spontaneous HLCx recordings for each subject to compare cortical state immediately before and between 10-30 minutes after SCI. Raw signals were down sampled to 500 Hz and then a Fast Fourier Transform (FFT) analysis was performed to confirm that the maximum peak frequency of the recordings was below 1 Hz. Frequency power (mV^2^) was extracted for each electrode by summing power within same frequency band: SWA (0.1-1 Hz), delta (δ, 1-4 Hz), theta (θ, 4-8 Hz), alfa (α, 8-12 Hz), beta (β, 12-25 Hz), low gamma (Lγ, 25-50 Hz) and high gamma (Hγ, 50-80 Hz). LFP power obtained from individual electrodes within a layer was averaged to obtain a layer frequency power. Finally, the power of each frequency band was normalized to the total power of the LFP recording (0.1-80 Hz) to obtain the relative power of each band. Additionally, individual up-states were selected within periods of 60 seconds of LFP for each subject to obtain the onset in control and acute SCI using the same methodology than for the onset of evoked-LFP responses (Fedchyshyn and Wang 2007). To calculate the velocity of propagation, the electrode presenting the earliest onset was taken as reference, and the velocity rate calculated as the distance in the cortical depth as a function of time.

### Thalamocortical recordings and analysis

Simultaneous electrophysiological recordings from FLCx and HLCx (n = 7 rats) in response to peripheral forelimb stimulation were obtained by using two single tungsten electrodes located on infragranular layer 5 of both cortical regions under control conditions and after SCI. Note that this dataset was obtained simultaneously to the thalamic data previously published in Alonso-Calviño et al. (2016), but the cortical dataset has been for the first time analysed for the present work. Anaesthesia and stimulation protocol were the same than for experiments explained above. Extracellular recordings were obtained using tungsten electrodes (TM31C40KT, 4-MΩ impedance at 1 kHz or TM31A50KT, 5-MΩ impedance at 1 kHz; World Precision Instruments Inc., Sarasota FL, USA). All recordings were pre-amplified in the DC mode, low pass filtered (<3 kHz) and amplified using a modular system (Neurolog, Digitimer Ltd). Analogue signals were converted into digital data at a 20 kHz with 16–bit quantization via a CED power 1401 apparatus (RRID:SCR_017282) controlled by Spike2. The data was analysed using Spike2 software. Onset latency of cortical responses was obtained using the same method than for S1HL multielectrode recordings phase (Fedchyshyn and Wang 2007) and evoked cortical responses to peripheral forelimb stimulation at 5 mA were used for this analysis.

### Histology

At the end of the experiments, animals were transcardially perfused with heparinised saline followed by 4% paraformaldehyde. Then the brain was removed and post-fixed in the same fixative solution for 24h at 4°C. After fixation, brain tissue was cryopreserved in a 30% sucrose solution until sank and coronal sections at 50 μm thick were obtained with a sliding microtome (Microm HM 450 V; Microm International GmbH, Germany). Following washing in 0.1 M phosphate buffer, sections were mounted in gelatin slides, air-dried, processed for tionine (Nissl) staining (T7029-5G, Sigma-Aldrich), dehydrated in xylene and coverslipped with DePeX (Cat#18243.01, SEVA, Heidelberg, Germany).

### Statistical analysis

Statistical analyses were performed using Statistica.Ink (RRID:SCR_014213, Statsoft Ibérica, Lisboa, Portugal). Shapiro-Wilk test was used to test normality distribution (p > 0.05). If normality was violated (non-gaussian distribution p < 0.05, rMUA responses magnitude, Fig 3), data were transformed to normal distribution by using square-root. Grubb’s test was used to identify and remove outliers (alpha = 0.05). Our main hypothesis (differences across layers) was first tested in our control data set (pre lesion) using one-way Analysis of Variance (ANOVA) with LAYER as independent factor. Dependent differences between pre- and post-injury among animals and layers were determined by two-way ANOVA, with LAYER as an independent factor and TIME as a repeated measures factor (two levels, PRE- and POST-lesion). When significant differences in analysis of variance where found, groups were further compared using Tukey’s *post hoc* test. For statistical analysis of the dual recording site from intact and deafferented cortices (Figure 6), we used two-tailed paired t-test with a *Bonferroni’s* correction for multiple comparisons (p < 0.025). The threshold for statistical significance was p < 0.05 throughout. Group measurements are expressed as mean ± standard deviation (SD) unless indicated in the figure legend. Results from statistical analysis are summarized in Table 1–3 of supplementary material. Graphs and figures were made using IgorPro (Wavemetrics, RRID:SCR_000325) and Adobe Illustrator (RRID: SCR_010279).

## RESULTS

### Laminar analysis of evoked neuronal responses in the somatosensory cortex following peripheral stimulation

We first characterized the evoked-local field potentials (evoked-LFP) across layers of the HLCx in response to sensory stimulation delivered either to the contralateral hindlimb (HL) or forelimb (FL) in intact anaesthetised animals (Fig. 1). For that, we used a linear 32-multielectrode probe inserted vertically into the HLCx such that recording sites were located across all layers in a single column to simultaneously record LFPs and local multiunit activity (MUA). Figure 1A shows a schematic representation of the recording location and stimulation paradigm as well as a histological preparation showing an electrode track into HLCx (Fig. 1B). Representative examples of averaged evoked-LFPs across the entire depth of HLCx in response to a 5 mA hindlimb stimulation (0.5 Hz) is shown in the left panel of Figure 1C. Note that high intensity stimulation was chosen to maximize the activation of peripheral nerves to ensure the full engagement of cortical circuits involved in somatosensory processing (Lilja et al., 2006). The laminar profile of averaged LFP responses of the population obtained by hindlimb stimulation showed a clear difference in the response magnitude across the cortical depth, with a maximum peak at distances from surface between 700-1000 μm corresponding to the thalamorecipient granular layer 4. No differences were observed using low intensity stimulation, which resembles light mechanical stimuli that preferentially activates dorsal column pathways (Lilja et al., 2006). Current source density (CSD) analysis was overlapped to LFP traces to determine the entrance of synaptic inputs in different cortical layers (Fig. 1C, heat map) showing an evident current sink (inward current) in L4. The active sink was surrounded by two strong current sources (outward current), one short-lasting in L2/3 and another long-lasting in infragranular layers (L5 and L6). A small and elongated current sink was also observed in L6, which may refer to the thalamocortical loop initiated by evoked responses. The greater LFP response magnitude and the current sink at the thalamorecipient L4 corroborate the correct location of the electrode at the somatotopic representation of hindlimbs. Finally, MUA responses (Fig. 1D, left panel) were analysed by rectifying the neuronal firing (rMUA) obtained in response to peripheral stimulation at high and low intensities (Fig. 1D, right panel). The area of the evoked-rMUA showed robust neuronal firing in the upper infragranular layer (i.e. L5), while L2/3 and L6 neurons showed low firing response. A similar laminar profile of neuronal responses was observed using low intensity stimulation (0.5 mA). The sparse and low firing observed in L2/3 under our experimental condition corroborates previous studies using distinct techniques such as *in vivo* and *in vitro* patch clamp, intracellular recordings (Wilent and Contreras, 2004; Jacob et al., 2017) and *in vivo* live Ca^2+^ imaging (Clancy et al, 2015).

We next examined the pattern of cortical responses when the peripheral stimulation was applied to an adjacent, non-corresponding somatotopic body region, i.e. recording in HLCx, while stimulating forelimb afferents. Figure 1E shows LFP representative traces produced in the HLCx in response to high intensity electrical stimulation of contralateral forelimb. Despite of the small amplitude, evoked-LFP responses were clearly observed across all recording channels with infragranular layers showing the highest magnitude. Similar results were also observed using 0.5 mA stimulation indicating a strong HL-FL cortical connectivity even to low intensity stimuli. In response to high stimulation, rMUA in infragranular layers showed a clear evoked activity with a peak in layer 6, while L2/3 and L4 exhibited only sparse and very low response. Therefore, our data indicate that each cortical layer exhibits a characteristic neuronal activity in response to stimulation of its own receptive field as shown in previous studies (Wilent et al., 2004; Sakata and Harris, 2009), but also in response to non-preferential peripheral stimulation (i.e. neuronal activity in HLCx in response to forelimb stimulation) which evidences a heterogenic cortical excitability most probably due to the composition of each local circuitry and synaptic inputs. Statistical analysis (one-way ANOVA, layer factor) comparing the response magnitude of cortical layers for each condition are summarised in supplementary table 1.

### Recording stability during the full transection of the spinal cord

We next sought to determine the effects that an immediate SCI had on the neuronal excitability of the deprived neurons across all cortical layers. As a first approach, we recorded the stability of our recordings during the exact moment of the spinal transection. Figure 2A shows original recordings of selected channels from an array of 32 electrodes inserted on the hindlimb cortex while applying 0.5 Hz electrical stimulation on contralateral hindlimb. Before SCI, all layers exhibited slow-wave oscillations composed of up and down-states and responded to evoked-sensory stimulation as evidenced by the fast and enlarged synaptic responses. While performing the recordings (and without removing the electrode from the hindlimb cortex), we proceeded with the full transection of the spinal cord. During the injury, the slow-wave activity is disrupted by only few seconds as observed by a short sustained depolarization observed across all layers of the deprived cortex (shaded area in Figure 2B). After this short period of time, cortical activity from hindlimb neurons returned to the previous slow-wave oscillations, while evoked-sensory responses were abolished, confirming the full transection of the spinal cord. The same pattern of short sustained depolarization was also observed simultaneous extracellular recordings obtained from layer 5 neurons of the intact, forelimb cortex and intracellular recordings electrophysiological recordings from layer 5 neurons in the hindlimb cortex (Figure 2C). The intracellular recordings of the layer 5 pyramidal neuron shows that the membrane potential and spontaneous activity were only disrupted by a short sustained depolarization with only few action potentials during the transection, and that the membrane potential immediately returned to baseline after this short period (Fig 2C, black traces, intracellular S1HL). In parallel, the extracellular forelimb cortical recordings (Fig 2C, green traces S1FL MUA) show parallel changes of activity, but it must be noted that the forelimb cortex is neither directly damaged nor sensory deprived by a thoracic SCI. The later confirm previous findings from our group showing that SCI affects brain states in both deprived and intact areas (Aguilar et al., 2010; Humanes-Valera et al., 2013). Therefore, these experiments demonstrated the neuronal stability of our recordings and the complete transection of the spinal cord.

**Figure 2:**
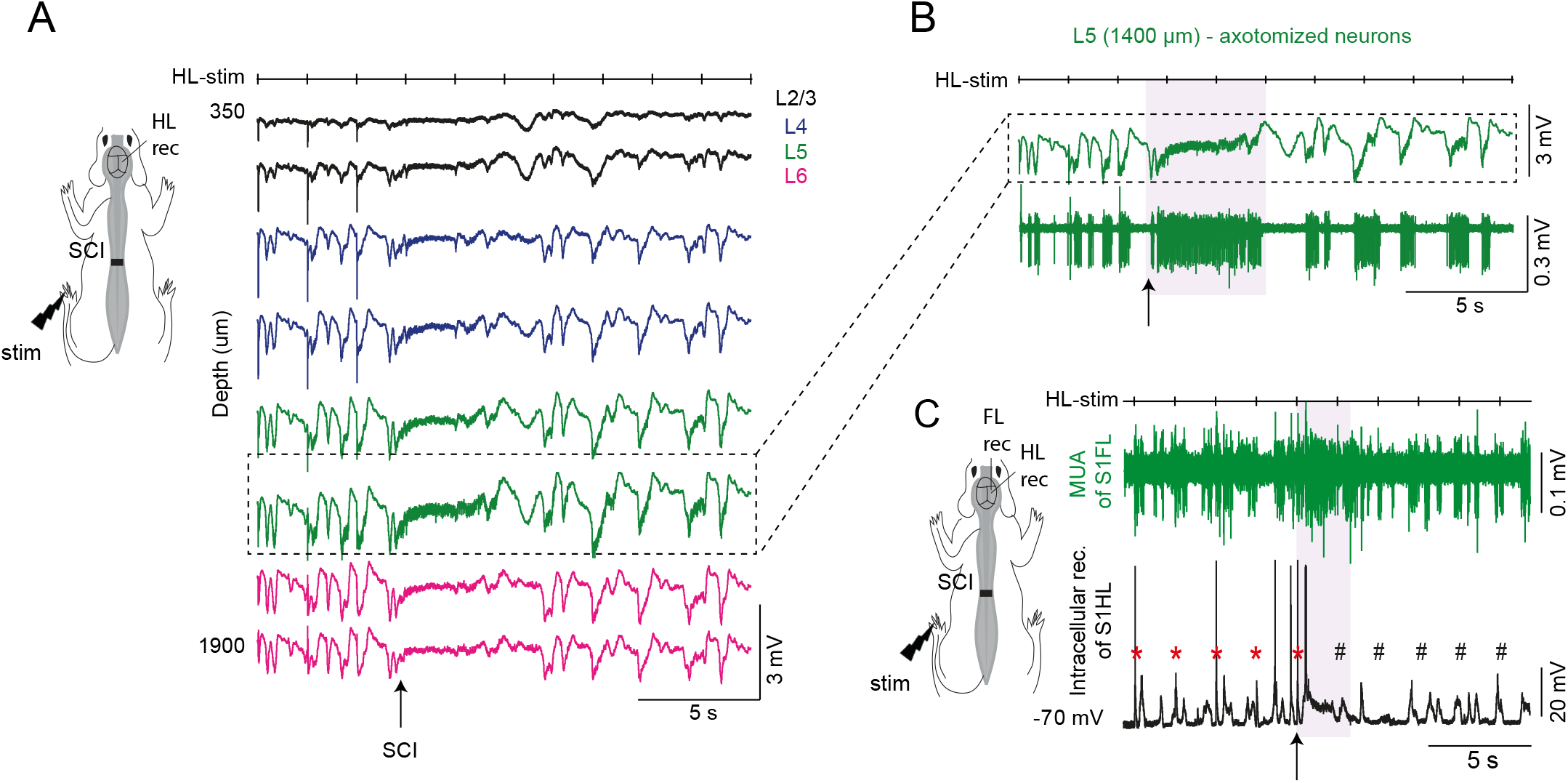
Stability of cortical recordings during spinal cord injury. (A) Original spontaneous LFP recordings obtained from distinct layers of hindlimb cortical location previous to SCI, during the moment of SCI (marked by an arrow) and immediately after SCI. Note that SCI triggers a transient (< 5s) increase of activity that could be associated to a short sustained depolarization. (B) Inset of the recording located in layer 5 (see identification by dotted rectangle in panel A). Extracellular action potentials observed after a high-pass filter (300-3000 Hz) evidenced that the immediate effect of SCI on cortical layer 5 is a transient (<5 s) and small increase of neuronal excitability (shaded area) that is re-synchronized after such short period to produce the characteristic oscillations of Up-states and Down-states of cortical slow-wave activity. Note in A and B the inclusion of upper channel named HL-Stim, indicating 0.5 Hz 5 mA hindlimb stimulation previous-during-post SCI in order to online confirmation of full spinal cord transection. (C) Simultaneous extracellular recording of layer 5 of intact forelimb somatosensory cortex (S1FL, green traces corresponding to high-filtered multiunit activity) and intracellular recording of a layer 5 neuron of S1HL (black traces). Arrow indicates the moment of full transection of the spinal cord. Note that neuronal depolarization (< 5 sec) after SCI is observed in both intact and deafferented cortices. Protocol of hindlimb stimulation (HL-stim, 0.5 Hz, 5 mA) is shown on top. Before SCI, HL-stim evoked action potentials and synaptic inputs in the intracellular recorded neuron (red asterisks) which was abolished after full transection of the spinal cord (black hashes).

### SCI induces layer-dependent functional changes in the sensory deprived cortex

Given the layer-specific activity induced in the HLCx in response to forelimb stimulation under control conditions, we hypothesised that SCI should induce specific effects on the evoked-LFP in a layer-dependent manner. To explore that, we investigated the neuronal activity across cortical layers immediately after sensory deprivation due to SCI in 24 rats (Fig. 3). As expected, an abolishment of the evoked-LFP was immediately observed in all layers of the HLCx in response to hindlimb stimulation, confirming the complete loss of sensory inputs (Fig. 3A). When peripheral stimulation was applied to the contralateral forelimb, the magnitude of evoked-LFP produced in HLCx were significantly increased in all layers but L6 (Fig. 3B, repeated measures ANOVA, SCI F_(1,91)_ = 50.2, p < 0.0001), indicating heterogeneous immediate changes in the excitatory post-synaptic activity of local pyramidal neurons and/or in the strength of arriving synaptic inputs. Next, we examined the area of the rectified MUA as a measure of local neuronal activity in each cortical layer recorded in a subset of SCI animals (16 out of 24 SCI animals; Fig. 3C). Figure 3D shows examples of evoked rMUA from each channel superimposed on a colour-coded map from the same animal in response to high intensity stimulation (5 mA) before and after SCI. Repeated measures two-way ANOVA of our data showed significant differences for layers (F_(3,60)_ = 12.5, p < 0.0001), and SCI condition (F_(1,60)_ = 18.1, p < 0.0001), but no interaction between layers and SCI (F_(3,59)_ = 1.2, p = 0.3145). Before SCI, L2/3 and L4 neurons exhibited very low activity, while robust firing was observed only in L5 and L6 in response to forelimb stimulation. After SCI, neuronal firing tended to increase in all layers (see original traces in Fig. 3E), with L5 rMUA significantly increased post-lesion (p = 0.0378). These results demonstrate that immediate sensory deprivation due to SCI produces increased neuronal firing in infragranular layers, and importantly creates differences in neuronal excitability among cortical layers of the deprived HLCx. Due to the non-consistent responses obtained among individuals when low intensity stimulation was applied, herein, data were considered only for responses obtained to high intensity stimulation in forelimb (5mA).

**Figure 3:**
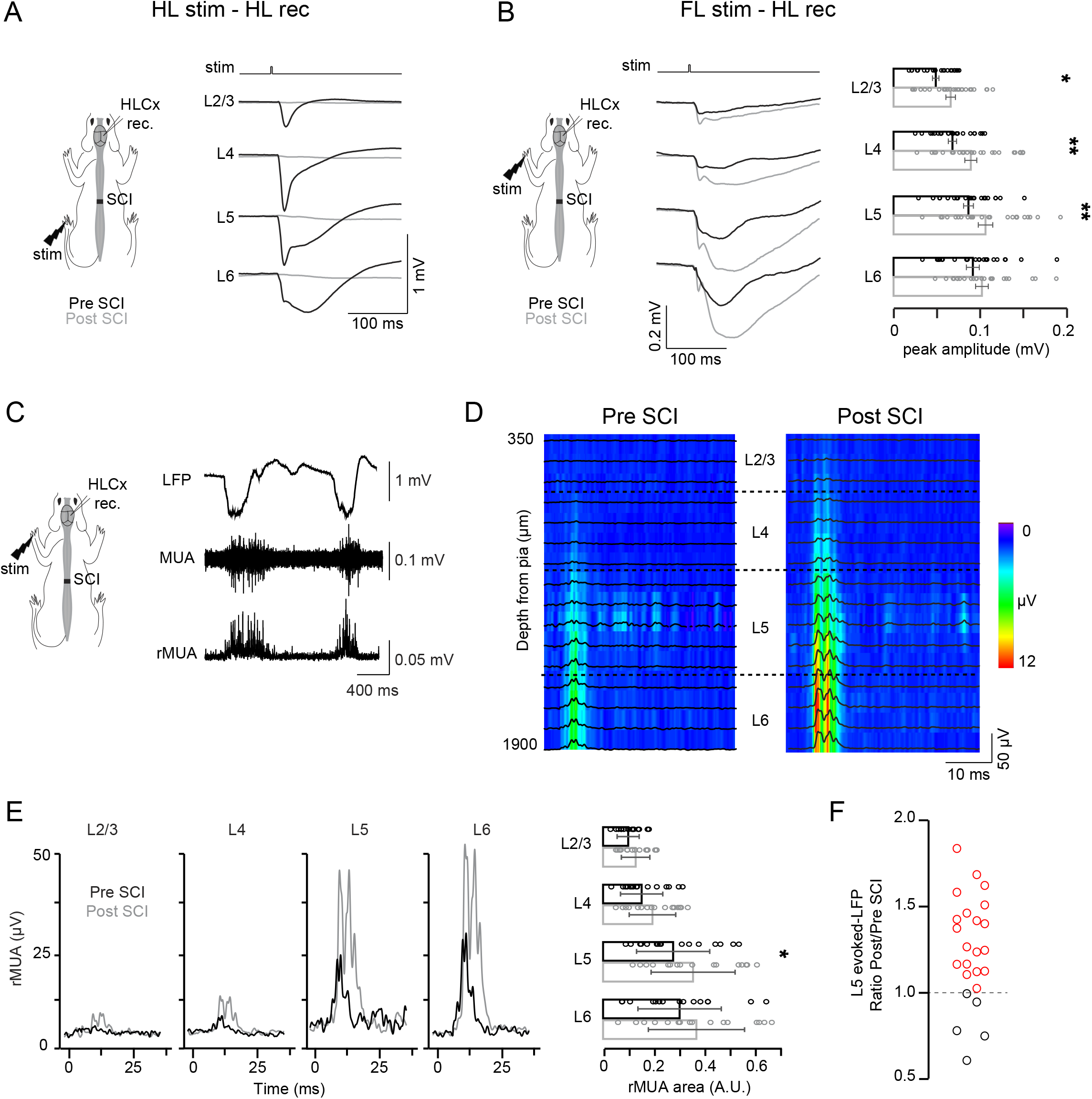
Spinal cord injury induces functional changes in the S1 deprived cortex in a layer-dependent manner. (A) Schematic showing the place of cortical recordings (HLCx rec) and hindlimb stimulation (HL stim). Examples of evoked-LFP averaged across electrode sites within each layer recorded in HLCx in response to HL stim (5 mA) in control (Pre SCI, black) and after SCI (Post SCI, grey). Note that complete spinal cord transection was confirmed by absence of evoked-LFP in HLCx. (B) Left, schematic showing HLCx rec and forelimb stimulation (FL stim) used for B-F. Examples of averaged evoked-LFP recorded in HLCx in response to FL stim (5 mA) in control and after SCI. Bars graph shows the population average of the evoked-LFP amplitude. (C) Original traces of LFP (top), multi-unit activity (MUA, middle) and rectified MUA (rMUA, bottom) in HLCx obtained following FL stim (5 mA). (D) Representative laminar profile of evoked-rMUA (black traces) overlapped to a colour map showing averaged HLCx responses to FL stim at 5 mA pre- and post-SCI. (E) Averaged evoked-rMUA across electrodes within distinct layers pre- and post-SCI and population averaged rMUA area. (F) Scattering plot showing ratio of layer 5 recordings calculated from evoked-LFP post-/pre-SCI. Dotted line represents ratio = 1. Dots represent individual experiments (red: ratio > 1, black: ratio < 1). Example traces in A-E are representative from the same animal. Data are mean ± SD (n = 24 rats for evoked-LFP and 16 rats for evoked-rMUA). * p < 0.05, ** p < 0.01. See also – Supplemental Table 2.

Previous data obtained from human studies have shown that changes in cortical activity are not homogenous across SCI patients (Wrigley et al., 2009; Freund et al., 2013; Jutzeler et al., 2015). Therefore, we hypothesised that immediate cortical deafferentation could affect non-uniformly to individuals in the studied population, and more specifically that a subset of our experimental animals may not exhibit the overall changes in the neuronal cortical activity following immediate SCI. To explore whether such assumption was true, we plotted the individual values corresponding to the ratios between the evoked-LFP response post and pre SCI obtained from layer 5 neurons for each studied animal (Fig. 3F). Note that layer 5 was chosen due to the large responses following forelimb stimulation and the significant effect previously described after SCI (Humanes-Valera et al., 2017; Fernández-López et al., 2019). This allowed us to identify a clear separation of animals in which SCI induced increments in the magnitude of the evoked-LFP (ratio > 1, 19/24 animals, 79 %), from those animals in which LFP amplitude remained stable or even decreased with a ratio < 1 (5/24 animals (21 %); Fig. 3F). Therefore, our results strongly suggest that SCI immediately increase the excitability of deprived neurons in a layer-specific manner. Moreover, we observed that immediate cortical deprivation does not affect homogeneously to all subjects of a population that could be clustered in affected and non-affected subsets.

### SCI strongly affects the corticocortical and thalamocortical connectivity between infragranular layers

Once we determined that SCI distinctly affected cortical layers and individuals, we sought a deep characterization of the functional alterations produced in the deprived HLCx in response to forelimb stimulation in both group of animals. It has been proposed that increased cortical responses after SCI could be mediated by a reduction of inhibitory activity that consequently unmask latent excitatory inputs of sensory deprived region, therefore modulating the dynamics of local responses (Sydekum et al., 2014). Taking this assumption into account, our proposed hypothesis was that sensory deprivation would differently affect the kinetics of LFP produced at each cortical layer based on the different inhibitory tone relieved at local neuronal network. Thus, we first analysed the slope values (decay rate in mV/s) of evoked-LFP that are often used to determine changes in the arrival and/or synchronization of synaptic inputs and are usually affected by changes in the excitation:inhibition balance (Fig. 4A). Under control conditions, hindlimb stimulation induced fastest responses in L4 of the HLCx (Fig. 4B), while forelimb stimulation induced similar slope values across all layers (Fig. 4C, black traces). After SCI, slope values significantly increased in HLCx layers 4-6 in response to forelimb stimulation in Group 1, but not in Group 2 animals (Fig. 4C). The fastest rising slope was observed in L6 indicating a strong recruitment of neuronal population after SCI as can be observed by a shorter time-to-peak values (Fig. 4D).

**Figure 4:**
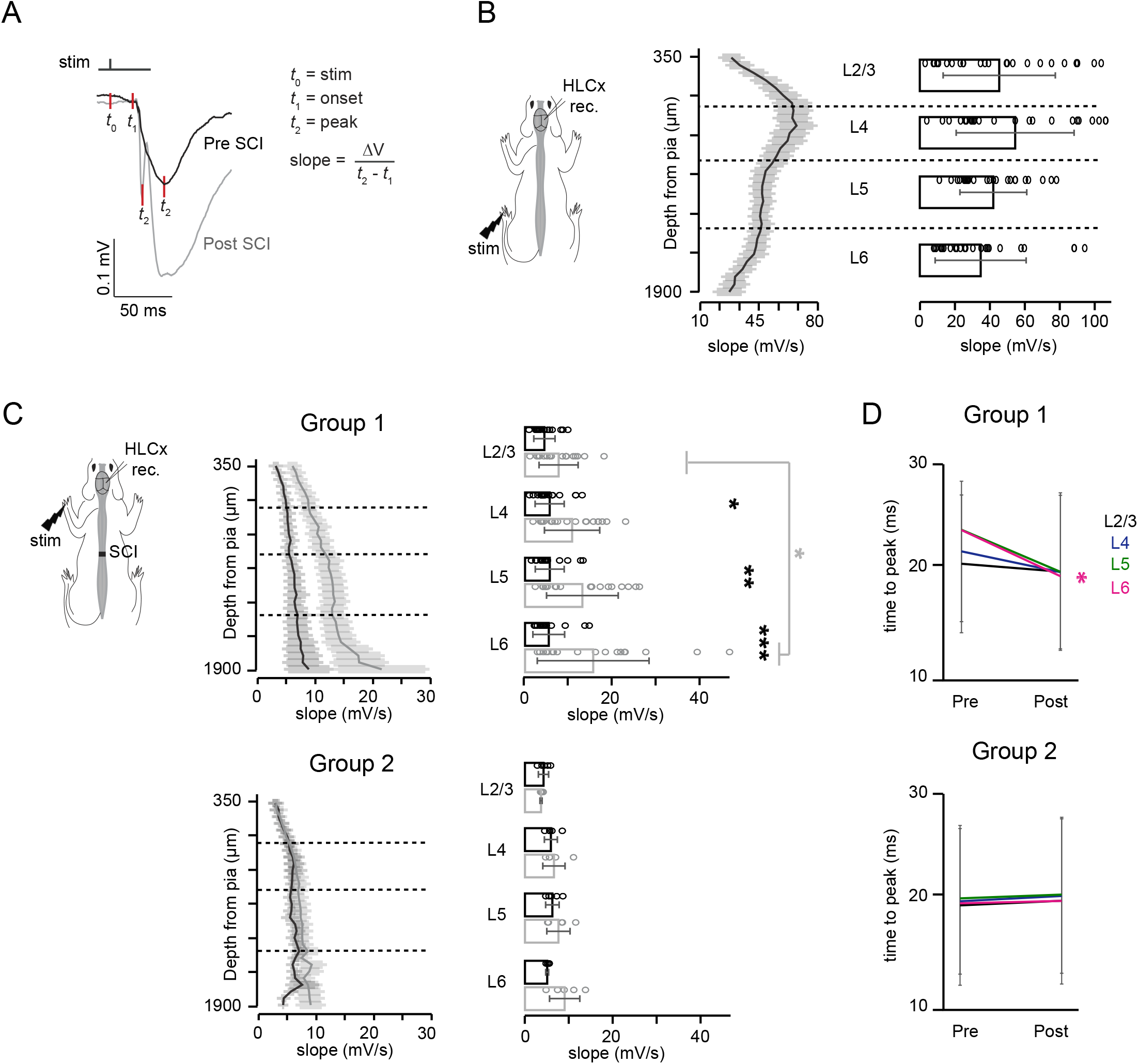
Laminar profile properties of evoked responses following SCI. (A) Original traces of averaged evoked-LFP in control condition (black) and after SCI (grey) illustrating the parameters used for slope analysis. (B) Scheme of an animal showing the place of cortical recordings (HLCx rec) and hindlimb stimulation (HL stim). Averaged slope of evoked-LFP from HLCx in response to HL stim at 5 mA in control. Bar graph shows the averaged slope values across electrodes within the same layer (n = 24 rats). (C) Averaged slope of evoked-LFP from HLCx in response to FL stim at 5 mA pre- and post-SCI for animals within Group 1 (top) and Group 2 (bottom). Bar graph shows the average slope values across electrodes within the same layer (n = 19, group 1, and n = 5, group 2). (D) Time-to-peak of the evoked-LFP responses before and after SCI from t0 to t2. Line graphs represent mean ± SEM only for illustrative purposes. Bar graphs data represent mean ± SD. * p < 0.05, ** p < 0.01, *** p <0.001.

Next, we determined the response latencies for each electrode site and for spatially averaged responses within electrodes from the same layer to create a spatiotemporal profile of the response onset (Fig. 5). Onset responses from LFP and rMUA were obtained by fitting a sigmoid function to the data and then computing the maximum curvature as described in Fedchyshyn and Wang, 2007 (see Methods). Figure 5A-B shows an example of laminar LFP profile and averaged onset latencies obtained from HLCx in response to hindlimb stimulation in control conditions. In this case, activity originated in upper middle layers and spread upwards and downwards resembling the propagation of evoked activity typically seen in distinct cortices (Sakata and Harris, 2009; Schroeder et al., 1998). Next, we analysed the evoked-LFP onset in response to forelimb stimulation, which could indicate how the corticocortical connectivity between HL-FL cortices is organized. The averaged onset of the animals was found to be similar across all layers with higher probability to be initiated in infragranular layers (Fig. 5C-E pre-SCI, black traces). These data indicate that population activity measured as LFP in response to peripheral forelimb stimulation reaches HLCx almost simultaneously as previously reported using voltage-sensitive dyes (Wester and Contreras, 2012). Such pattern of activity onset was strongly affected by sensory deprivation in a layer-dependent manner. Infragranular layers presented a significant delay in the onset of the evoked responses, while onsets in granular and supragranular layers were not affected (Fig. 5C-E post SCI). In addition, onset probability was also similar across layers after SCI (26-32% for L4-L6 and 16% for L2/3). In this case, both groups of animals showed similar functional changes for onset-latency responses (Fig 5D group 1, Fig 5E group 2). We then analysed the onset pattern of the evoked rMUA (Fig. 5F). Before SCI, HLCx infragranular layers of group 1 exhibited faster onset latencies than supragranular layer with neuronal firing originating mostly in infragranular 6. After SCI, differences in onset responses disappeared due to rMUA of infragranular tend to be delayed (Fig. 5F, left graph). Group 2 showed high variability due to the small sample (n = 3; Fig 5F, right graph).

**Figure 5:**
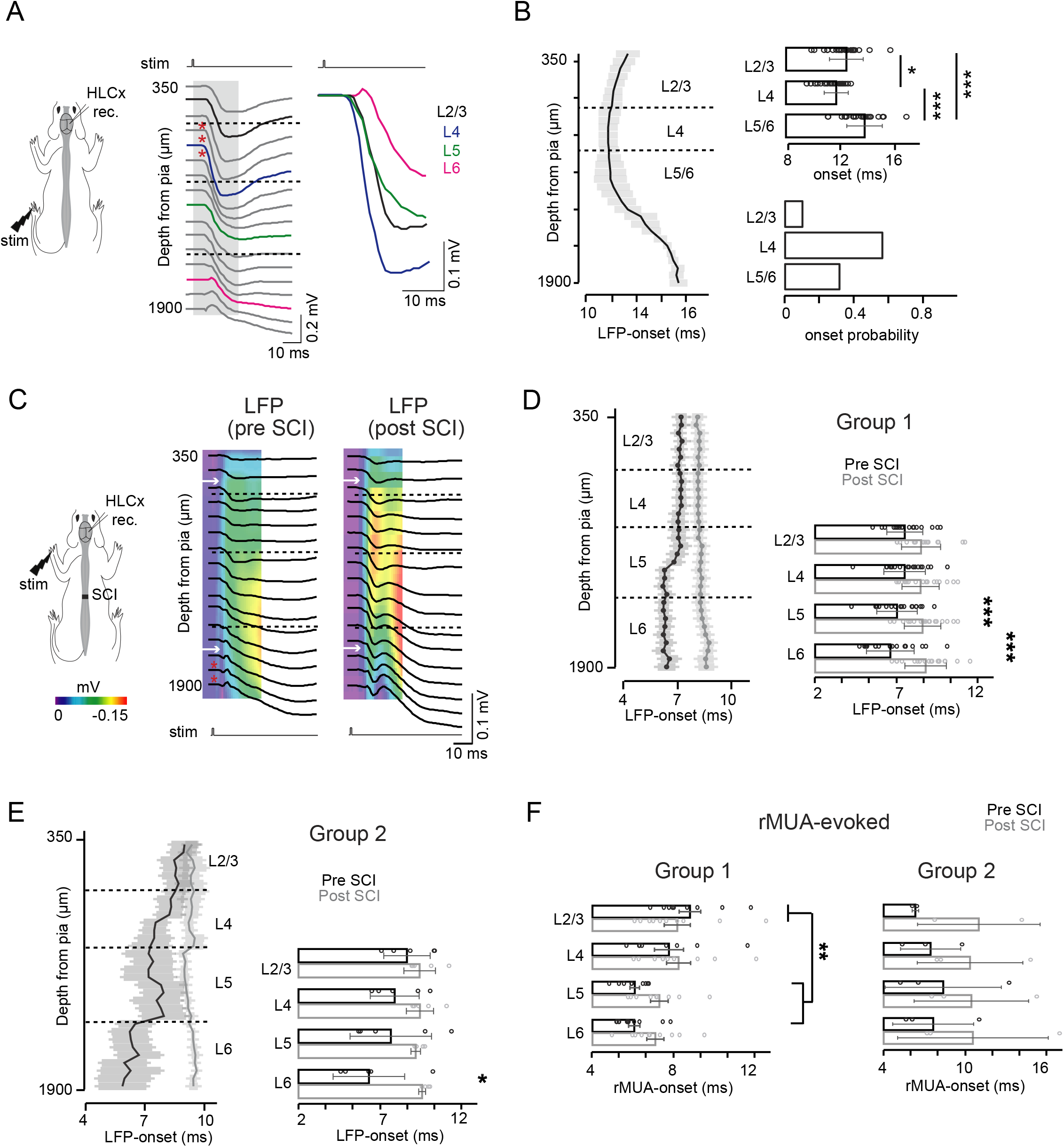
Deprived infragranular layers exhibits delayed evoked-onset responses. (A) Left: onset profile of evoked-LFP in HLCx in response to hindlimb stimulation (HL stim) before SCI. Red markers indicate electrodes in which evoked response started. Grey area indicates the period of response onset taken to highlighted LFP traces (in black) in the right. (B) Latency-onset averaged profile to hindlimb stimulation (5 mA) at left. Note the onset in the thalamorecipient granular layers (L4). Right top: bar graph of the evoked-LFP onset responses. Right bottom: normalized histogram of the evoked-LFP onset probability from. Data obtained from 24 rats. (C) Onset profiles from the same animal pre- and post-SCI showing averaged evoked-LFP in HLCx in response to forelimb stimulation represented as in A, overlapped on colour maps from the same traces. White arrows indicate the onset response in L2/3 and L6. In D-E, latency-onset of the laminar profile (at left) and averaged across layers (at right) by forelimb stimulation (5 mA) is represented for Group 1 (D) and Group 2 (E) for evoked-LFP analysis. In (F), response latency-onset was quantified for rMUA-evoked of group 1 (left) and group 2 (right). Dots represent individual experiments. Line graphs represent mean ± SEM only for illustrative purposes. Bar graphs data represent mean ± SD (n = 19 rats for group 1 and n = 5 for group 2). * p < 0.05, *** p < 0.001.

### SCI induces delayed thalamocortical inputs to the deprived cortex

A possible mechanism leading to the delayed LFP and rMUA onsets in infragranular layers would be that SCI induces changes in the two main neuronal pathways that drive the peripheral information from forelimbs to HLCx: 1) a canonical corticocortical pathway in which synaptic inputs from the thalamic forelimb region routes to FLCx and then reaches HLCx mostly through L2/3 (FL-Th➔FLCx➔HLCx); and 2) a non-canonical thalamocortical pathway involving the activation of a subset of HL neuronal population in the thalamus that project to HLCx in response to forelimb stimulation, HL-Th➔HLCx (Figure 6A; Alonso-Calviño et al., 2016; Francis et al., 2008). Considering that delayed onset was observed in both HL and FL thalamus immediately after SCI (Alonso-Calviño et al., 2016), we wanted to determine which thalamocortical pathway was the most plausible to be involved in the delayed HLCx evoked-responses. To prove this idea, we have analysed simultaneous electrophysiological recordings from FLCx and HLCx obtained from tungsten electrodes located on layer 5 of both cortical regions under control conditions and after SCI (Fig. 6B, these data were obtained simultaneously to thalamic data that were published in Alonso-Calviño et al., 2016). We postulated that if increased response latencies of HLCx after SCI were induced by changes in the canonical pathway, then synaptic inputs would arrive earlier at the FLCx than at HLCx. On the other hand, if non-canonical pathway produces the longer latency of HLCx evoked responses, and then both cortical regions should exhibit similar latencies after forelimb stimulation. Our results demonstrated that both the intact FLCx and the sensory deprived HLCx showed similar increased latencies to peripheral FL stimulation immediately after SCI with no differences between them (Fig. 6C, pre-SCI p = 0.2190 and post-SCI p = 0.8889). Therefore, increased latency of cortical evoked responses could be at least in part due to longer latencies that take place in thalamic VPL corresponding to HL and FL as we described in Alonso-Calviño et al. 2016. Moreover, we cannot discard other intrinsic properties or corticocortical mechanisms involved in this process.

**Figure 6:**
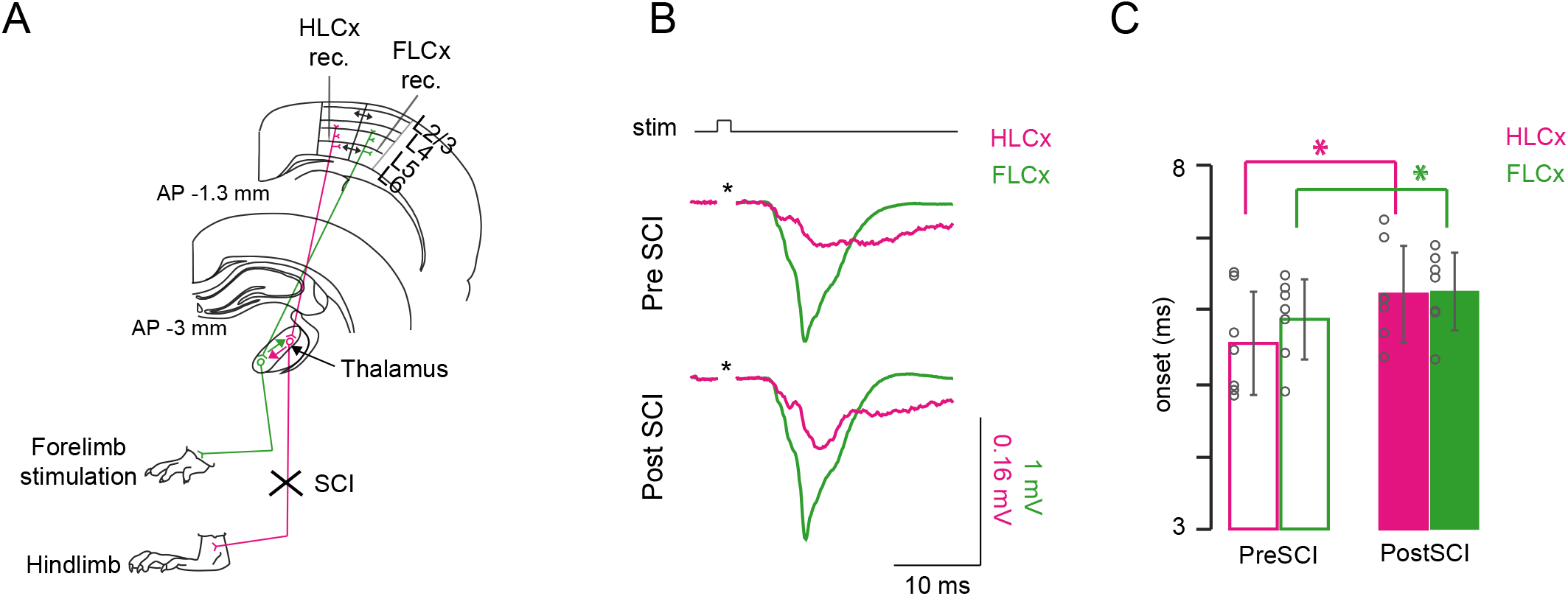
Thalamocortical responses in HL cortex are delayed after SCI. (A) Scheme showing the thalamocortical pathways activated after forelimb and hindlimb stimulation. Extracellular recordings were simultaneously obtained from L5 of HLCx and FLCx while forelimb stimulation at 5 mA was applied. Colored arrows in thalamic nucleus represent the collaterals allowing reciprocal activation of neighbor regions. (B) Original field potential traces obtained from simultaneous recordings from HLCx (magenta traces) and FLCx (green traces) before and after SCI. (C) Averaged bar graphs displaying onset latencies of evoked responses in HLCx and FLCx. Data are mean ± SD (n = 7 rats). * p < 0.025.

### Spontaneous activity is determined by cortical layer properties under control conditions and after sensory deprivation

Cortical spontaneous activity in anesthetized animals is generally dominated by slow-wave activity (SWA, Fig 7A). SWA is mainly originated from local neuronal networks and corticocortical connections (Chauvette et al., 2010; Sanchez-Vives and McCormick, 2000) and is importantly modulated by sensory inputs reaching the cortex throughout thalamic pathways (Rigas and Castro-Alamancos, 2009). In this context, we have previously demonstrated that the drastic sensory loss after SCI reduces the neuronal excitability during cortical SWA (Aguilar et al., 2010; Fernández-López et al., 2019). As these data were obtained from deprived HLCx layer 5 neurons and SWA up-states propagate vertically across layers *in vitro* and *in vivo* (Sakata and Harris, 2009; Sanchez-Vives and McCormick, 2000), we thus calculated the onset of the spontaneous activity to determine the pattern of vertical propagation in our experimental conditions from Group 1 animals. Before SCI, spontaneous up-states initiated in any cortical layers (Fig. 7B), with infragranular layers 5 and 6 showing the highest onset probability (~80%), which corroborates previous findings in somatosensory cortex of anesthetized rats (Fiáth et al., 2016; Sakata and Harris, 2009). After SCI, we found that although spontaneous up-states also tend to start in infragranular layers, this probability decreased to ~60%. This effect was achieved by a parallel increase in the probability of up-states starting at L2/3 (Fig 7B). By analysing the rate of neuronal activity transfer across layers, we observed that up-states originated either in L2/3 (Fig. 7C-D) or L5/6 (Fig 7E-F) propagated downwards and upwards, respectively, in similar velocities before and after SCI. Therefore, while the neuronal network of the deprived cortical column responsible to transfer sensory information along cortical depth is not immediately affected by SCI, the increased probability of spontaneous up-states initiation in L2/3 indicates a plausible enhancement of corticocortical connections between HL-FL that helps to propagate spontaneous activity from the adjacent FL cortex.

**Figure 7:**
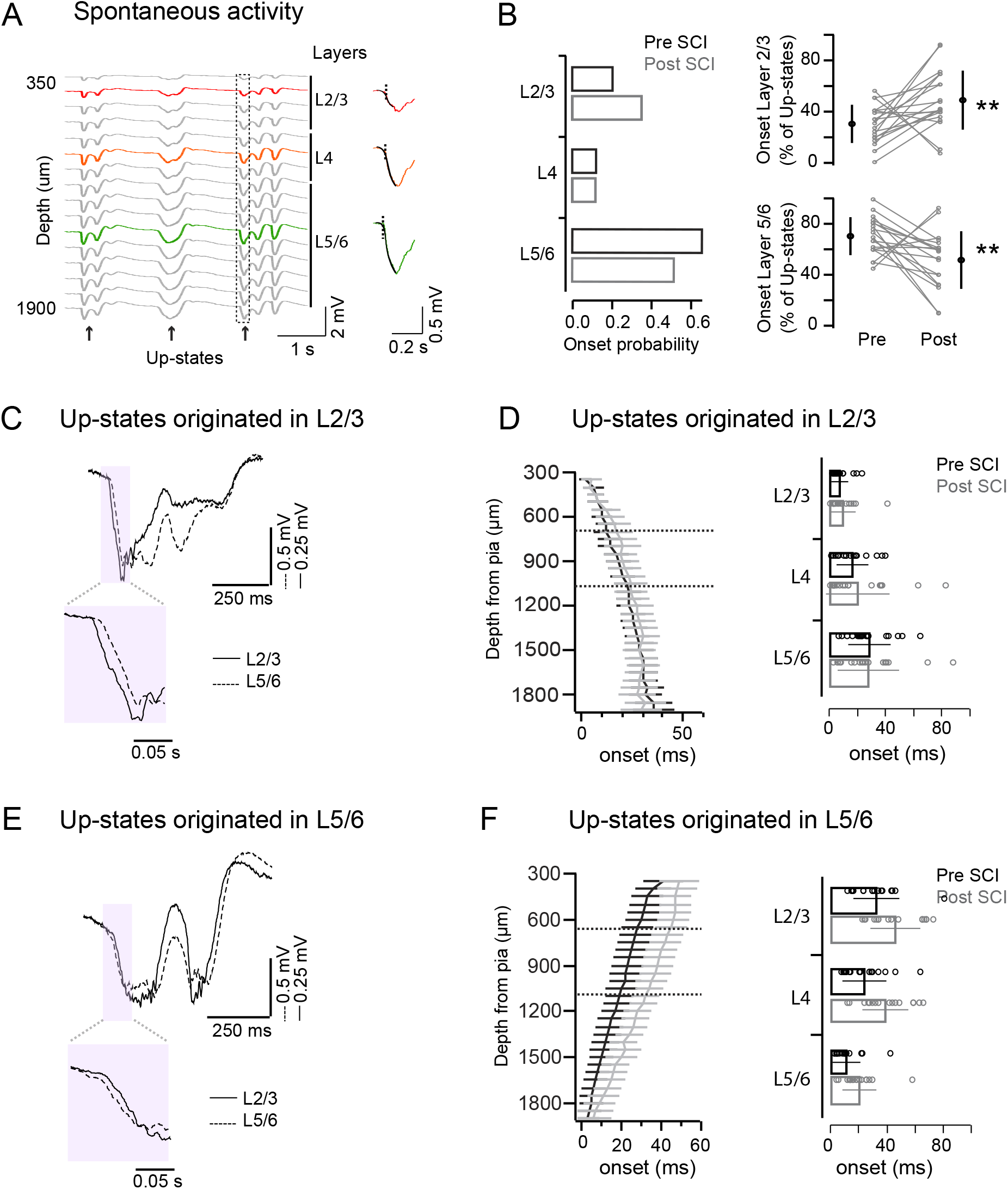
Onset and propagation of spontaneous up-states is altered after SCI. (A) Example SWA across electrodes (left) showing insets of the sigmoid fitting used to calculate the onset latency (right). Colors indicate distinct layers. (B) Left: Histogram of onset probability across layers of spontaneous up-states pre- and post-SCI. Right: Percentage of up-states with origin in layer 2/3 or layer 5/6 pre- and post-SCI. Lines represent each individual. Black dots are mean ± SD. (C) Representative example of spontaneous up-state with origin in layer 2/3 (black line) and delayed L5/6 (dotted line). Insets are magnification of magenta areas. (D) Laminar profile and averaged onset latencies of up-states originating in layer 2/3. (E) Representative example of spontaneous up-state with origin in layer 5/6 (dotted line) and delayed L2/3 (black line). (F) Laminar profile and averaged onset latencies of up-states originating in layer 5/6. Data are mean ± SD (n = 19 rats). * p < 0.05, ** p < 0.01.

Changes in the generation of spontaneous activity may indicate that other cortical features could be affected by SCI. Following this idea, we considered that alterations of intrinsic excitability should also be reflected in the frequencies content of the LFP signal in each layer during spontaneous activity from Group 1 animals (Fig. 8). Spectrogram analysis performed on individual traces from L2/3, L4 and L5/6 layers (exampled red traces on Fig. 8A) confirmed the presence of slow (0.1-9 Hz) and fast rhythms (10-80 Hz) during up-states before SCI (Fig 8B). Interestingly, our spectrogram analysis showed an increased relative power of fast rhythms mainly in supragranular layers following immediate SCI. Changes in the internal frequencies within up-states after SCI can be clearly observed by the fast oscillations (band pass filtered 25-80Hz) shown in the original traces in Figure 9C (lower traces). To explore such frequency differences in a systematic manner, we performed a power spectrum analysis (using Fast Fourier Transform, see methods) of the LFP frequencies content that were divided into different bands: SWA (0.1-1 Hz), delta (1-4 Hz), theta (4-8 Hz), alpha (8-12 Hz), beta (12-25 Hz), low gamma (25-50 Hz) and high gamma (50-80 Hz). Our data showed that from all the studied frequency bands, the relative power of the high gamma was consistently increased in supragranular layers following a SCI (Fig. 8D) while SWA, that characterized the cortical state imposed by the anaesthesia, did not changed. The rest of the studied frequency bands were not altered by SCI as summarized in Supplemental Table 3. Therefore, a massive sensory loss produced by an immediate SCI affects differently to of intrinsic excitability of cortical layers characterized by lower ability of layer 5 to generate up-states and increased gamma frequency in supragranular layers during spontaneous activity.

**Figure 8:**
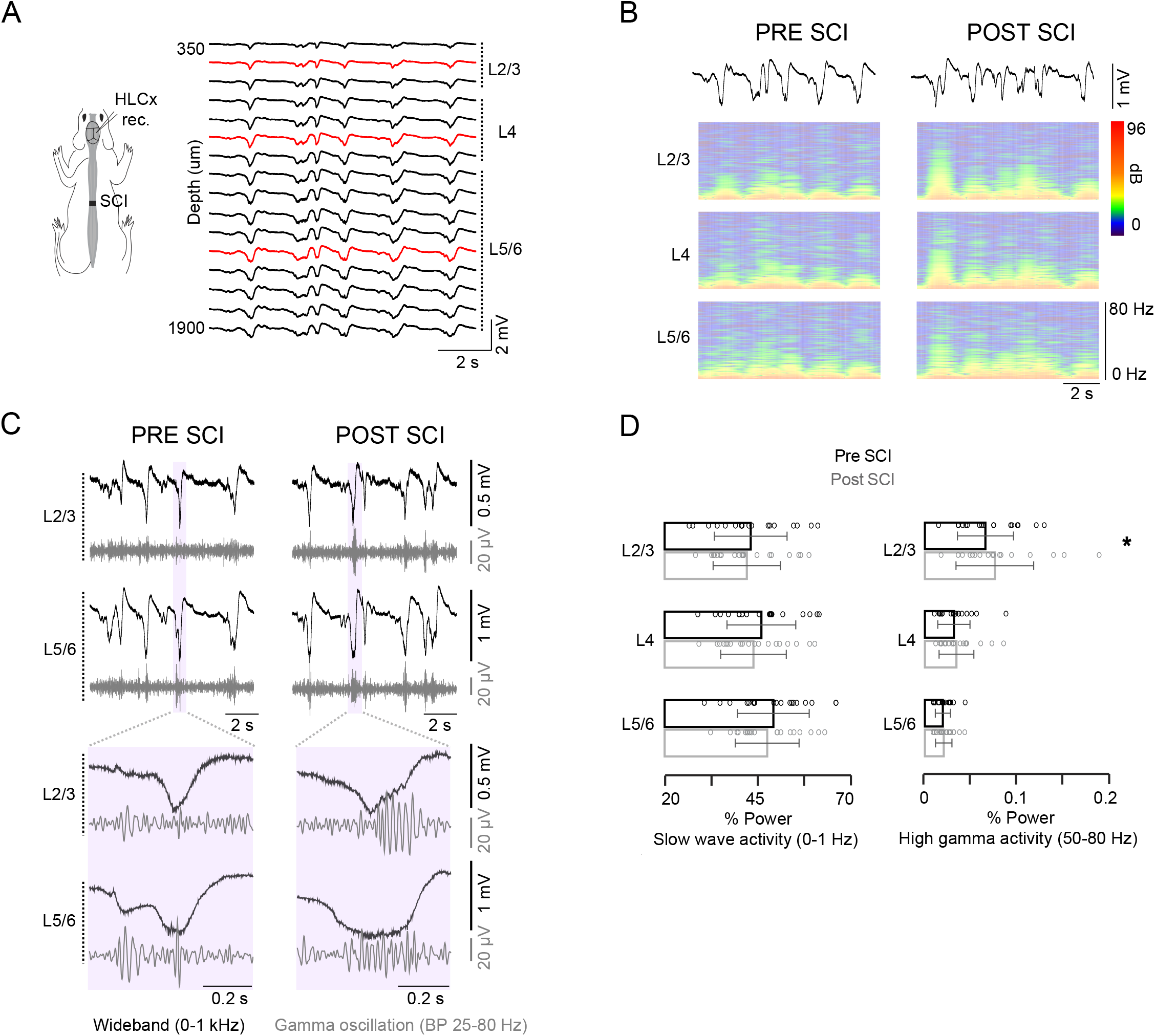
SCI increases high-gamma frequency oscillations in supragranular layers. (A) Example of spontaneous SWA profile recorded from HLCx in control conditions. Red traces represent selected electrodes within each cortical layer used for B-C. (B) Color plots showing spontaneous SWA spectrogram in L2/3, L4 and L5/6 pre- and post-SCI. (C) Wideband signal (black) and gamma-filtered (25-80 Hz, grey) traces of SWA from L2/3 and L5/6 with expanded traces of the indicated time window. (D) SWA and gamma relative power from distinct layers before and after SCI. Data are mean ± SD (n = 19 rats; ** p < 0.01). See also – Supplemental Table 3.

## DISCUSSION

Here we investigated the immediate effects that a robust sensory deprivation induces in distinct layers of the primary somatosensory cortex. Our study gives strong evidence that acute SCI induces layer-dependent changes in local circuits mediating evoked and spontaneous activity in the deprived cortical region through alterations of both corticocortical and thalamocortical connections. Regarding evoked responses, sensory deprivation potentiated the response magnitude and the rising of the population neuronal activity of infragranular layers (L5/6) of the deprived HLCx. On the other hand, the study of spontaneous activity show that supragranular layer 2/3 is the most affected by SCI exhibiting increased probability to initiate spontaneous up-states and increased power of high-frequency oscillations in the gamma band spectrum. Therefore, our results show that local neuronal and network properties of each cortical layer are responsible for differential effects observed in the deprived somatosensory cortex after SCI.

### Layer-dependent changes in sensory-evoked responses after SCI

Sensory deprivation has dramatic effects on the organization of brain circuitries, leading to a takeover of the deprived cortex by other cortical areas. This process is initiated as soon as deprivation occurs (Humanes-Valera et al., 2013) and continues in a time scale from days-to-months (Endo et al., 2007; Sydekum et al., 2014; Humanes-Valera et al., 2017; Fernández-López et al., 2019). In the case of SCI, the reorganization of the deprived cortex leads to the acquisition of new sensory functions that could help functional recovery (Rossignol and Frigon, 2011) as well as initiates associated pathologies as pain and spasticity (Siddall and Loeser, 2001). The mechanisms driving beneficial or detrimental reorganization are unknown, but it could rely on the complex laminar organization of cortical areas known to have distinct cellular composition and intrinsic circuitries. Therefore, a better knowledge of the contribution of each cortical layer to the well-known phenomenon of CoRe after sensory deprivation is required. In this study, we have included for the first time the perspective of cortical layering role in the cortical changes after SCI, which could explain initiation and complexity of CoRe as well as explain the variability between individuals as observed in human patients.

Previous differences among layers were only addressed in a neonatal SCI model in which the effects of exercise in the cortical long-term plasticity were studied (Kao et al., 2009). Our present data goes further to demonstrate that neuronal activity is differentially affected across layers of the sensory-deprived HLCx immediately after a SCI in adult individuals. Under our experimental conditions, increased magnitude of evoked-LFP in response to stimulation of the contralateral forelimb was observed across layers of the deprived cortex (Fig 2B), which is very consistent with results obtained using brain scanning approaches (Endo et al., 2007; Ghosh et al., 2010). This effect could be explained by the anatomical and functional overlapping of hindlimb and forelimb cortical areas (Moxon et al., 2008; Morales-Botello et al., 2012), in which corticocortical excitatory inputs may became unmasked following SCI and increase the responses to stimulation of a non-corresponding extremity (i.e. forelimb). In addition, we have previously shown that SCI increases neuronal responses in the thalamic hindlimb region to forelimb stimulation (Alonso-Calviño et al., 2016). Therefore, changes in thalamic excitability could also play a direct role in the increased cortical responses of sensory deprived HLCx. The initial slopes of evoked-LFP, often used to determine changes in the arrival and/or synchronization of synaptic inputs and are usually affected by changes in the excitation:inhibition balance, were significantly faster in granular and infragranular neurons (Fig 4C). Since cortical layers are known to display different inhibitory features (Wilent and Contreras, 2004), our data showing changes in the LFP slope in a layer-dependent manner suggest that local network properties are differentially affected by SCI and could represent an unequal reduction of local inhibition across layers allowing infragranular cells to better integrate evoked-sensory inputs.

Onsets of evoked-sensory responses in cortical regions are driven by synaptic inputs from corticocortical and thalamocortical connections. Our data showing almost simultaneous initiation of evoked responses between layers (Fig. 5C-D) corroborates *in vitro* data showing that horizontal corticocortical connectivity with adjacent cortical areas induces similar onsets of evoked responses (Wester and Contreras, 2012). However, this feature was strongly delayed in L5/6 of the deprived cortex after SCI, which could indicate modifications in the cortico-cortical synaptic connectivity but also in thalamic connections that project to HLCx. *In vivo* peripheral forelimb stimulation induces strong neuronal responses in the FL area of the thalamic VPL, but also in a population of the HL area of the VPL through collaterals that finally project onto HLCx (Fig. 6A, Alonso-Calviño et al., 2016). By simultaneously recording neuronal activity in HLCx and intact FLCx we showed that SCI leads to a delayed latency in the neuronal activity of both regions in response to peripheral stimulation. In addition, we did not observe changes in the onset of thalamus-devoid supragranular neurons after SCI (Fig 6B), suggesting that the onset of evoked-LFP responses in infragranular layers is more conditioned by changes in thalamic inputs. Therefore, our data indicate that although a delay in the arrival of synaptic inputs in infragranular layers is observed after SCI, these are more efficiently integrated as showed by increased slope and magnitude of evoked-LFP. Then, sensory deprivation produces changes in the integration properties of local networks of the deprived infragranular neurons that could be the basis for the long-term reorganization of sensory cortex observed after SCI.

Electrophysiological recordings of LFP and action potentials reflect different but complementary aspects of neuronal processing. While LFP integrates subthreshold activity as synaptic inputs and membrane potentials from a neuronal population in a given region, action potentials are the output signals from individual neurons close to the recording electrode. Contrary to the homogeneous increase in the evoked-LFP after SCI through L2/3-L5, we found striking differences among layers regarding MUA. In this case, sensory deprivation increased neuronal excitability in infragranular neurons of the deprived HLCx but not in granular and supragranular layers. Infragranular neurons have several characteristics that may lead to most of the changes: they receive excitatory inputs from all other cortical layers and neighbouring cortical areas (Schubert et al., 2007), they receive extensive thalamic inputs, and they have larger receptive fields (Moxon et al., 2008; Rigas and Castro-Alamancos, 2009; Wester and Contreras, 2013). Therefore, changes in the local network connectivity and/or intrinsic properties of infragranular neurons are more prone to be noticed after SCI compared to other layers. Moreover, the only intracellular data obtained from infragranular neurons after acute SCI (Humanes-Valera et al., 2017) perfectly support the relation between the increased MUA responses and the faster slope of evoked-LFP that we show in the present work. Regarding the changes in layer 5 neurons, we would like to remark that although previous data have shown preservation of L5 pyramidal neurons after axotomy due to SCI (Ghosh et al., 2012), we cannot discard that direct damage at the level of spinal cord of corticospinal neurons with origin in cortical layer 5 may have influence in some of the observed functional changes. Therefore, our data strongly indicate that excitability of infragranular neuronal networks are mostly affected in the context of evoked responses, which could be directly implicated in the mechanisms regulating subcortical output generating adaptive behaviour and functional recovery following spinal cord injury.

### Sensory deprivation affects the generation of gamma oscillations and the propagation of Up-states in the cortical column

In our experimental model, the cortical activity before SCI was settle to the state of slow-wave oscillation (~1 Hz) characterized by alternating periods of synchronized activation of neuronal population (up-states) and silent periods (down-states; Sanchez-Vives and McCormick, 2000). Up-state events are dominated by gamma frequency activity (25-80 Hz) with implications in multiple aspects of information processing such as sensory representation (Castro-Alamancos, 2009), sensorimotor integration (Schoffelen et al., 2011) and cognition (Gruber et al., 2004). Here, we describe that sensory deprivation due to SCI induces a layer-specific modulation of high frequency oscillations during Up-states, with gamma range being strikingly increased in supragranular layers, but not in infragranular or granular layers. Layer 2/3 is known to present a network of inhibitory neurons that initiates gamma-oscillations either through PV-neurons (Cardin et al., 2009; Welle and Contreras, 2016) or somatostatin neurons (Veit et al., 2017). In this context, the sensory deprivation produces a reduction in the constant thalamic excitatory inputs onto the deprived cortex that may lead to an increase of the general inhibitory tone in supragranular layers which facilitates local mechanisms of gamma oscillations. In addition to the well-described role of high frequencies in information processing, gamma oscillations have also been implicated in the formation of cortical maps during development (Minlebaev et al., 2011) and related to generation of pain perception in somatosensory cortex (Tan et al., 2019). Therefore, it is plausible that the increased gamma may be related with several long-term physiological changes observed following SCI such as cortical reorganization linked to functional recovery as well as maladaptive plasticity linked to chronic pain. Moreover, this neuronal feature could also be used as a functional biomarker for CoRe after a CNS injury, as previously shown in human auditory cortex after noise trauma (Ortman et al., 2010).

Spontaneous up-states within slow-wave oscillations usually initiate in deep infragranular layers (Sakata and Harris, 2009) and depend primarily on both intrinsic properties of the cortical column and corticocortical connections (Sanchez-Vives and McCormick, 2000). Our data shows that SCI induces a 1-fold increase in the probability of up-states generation in layer 2/3 with a consequent decrease in layers 5/6. There are several possible mechanisms that both isolated or synergistically could be leading to such changes. First, subthreshold oscillations in the membrane potential during gamma oscillations facilitate the generation of spontaneous up-states (Puig et al., 2008). Since we also observed increased gamma in L2/3 after SCI probably due to increased activity of inhibitory neurons, this mechanism could be a trigger to the increased up-state onset (Compte et al., 2003). Second, we have previously shown that SCI decreases neuronal activity during up-states in layer 5 neurons (Fernandez-Lopez et al., 2019), which may favour the initiation of up-states in layer 2/3 by releasing supragranular neurons from L5 to L2/3 modulation (Wester and Contreras, 2012). In the same way, *in vitro* experiments show that up-states generation in the somatosensory cortex is reduced after blocking thalamocortical inputs, while a reduction of the fast excitatory activity (mediated by AMPA receptors) increases the probability to generate up-states in cortical layers 2/3 (Favero and Castro-Alamancos, 2013). Finally, the superficial cortical layer 2/3 is also known to receive long-range axons from other cortical areas that facilitate long-range synchronization (Yamashita et al., 2018). We have previously shown that SCI induces neuronal changes not only in the deprived HL cortex, but also in the adjacent, intact forelimb cortex (Humanes-Valera et al., 2017; Humanes-Valera et al., 2013; Yague et al., 2014). Therefore, changes in oscillatory synchronization in the sensory forelimb cortex may be transfer to hindlimb cortex through L2/3 corticocortical connections during spontaneous activity facilitating the up-state generation within the deprived cortical column. Despite of the mechanism used by the deprived cortex to generate spontaneous activity and to propagate the neuronal information across layers, the increase in L2/3 up-states may allow the deprived column to maintain its internal activity with possible implications in the processing of evoked sensory inputs as well as the reorganization of cortical areas after sensory deprivation.

Taking our results in the perspective of the long-term physiological effects that a SCI produces in the primary somatosensory cortex, it has been described how functional changes known as cortical reorganization can benefit functional recovery or induce pathophysiological consequences (i.e. neuropathic pain). There are two main factors involved in the development of CoRe: 1) structural changes of neuronal networks and connections linked to anatomical rewiring of axons and dendrites, and 2) functional changes linked to activity-dependent plasticity and/or homeostatic plasticity (Muret and Makin, 2020). Since our results were obtained in a narrow time window (from minutes to few hours after deprivation), the possibility of structural and/or anatomical changes is limited. On the contrary, we consider that the observed neuronal network alterations in a specific layer will depend on the importance of lacking the preferred connectivity (thalamocortical or corticocortical), the inhibitory neuronal composition and how the network integrates secondary/non-preferential and weak connections (as described in Muret and Makin, 2020). Then, immediate functional changes described in our results point to an initiation of homeostatic processes intended to compensate input deprivation by rebalancing excitation:inhibition as it has been described in other sensory systems (Keck et al., 2013; Teichert et al., 2017), which can be followed in the long-term by a process of activity-dependent plasticity. We found our results consistent with other deprivation models as amputation (Makin et al., 2013), in which has been described that the map remains stable, but the function is affected. Importantly, our work provides a new framework for a better understanding of CoRe after SCI by identifying a role for deprived supragranular layers in better integrating spontaneous corticocortical information to modify the excitability of the column, and for deprived infragranular layers in better integrating evoked-sensory inputs to preserve specific corticothalamic and cortico-subcortical networks. We postulate that the layer-specific neuronal changes observed immediately after sensory deprivation may guide the long-term alterations in neuronal excitability and plasticity linked to the rearrangements of somatosensory networks and the appearance of central sensory pathologies usually associated with SCI.

### Preclinical relevance of the findings

Our data also identified a subset of animals (21%) that consistently didn’t show immediate neuronal changes related to neither evoked responses nor spontaneous activity. Interestingly, population studies comparing neuronal activation in SCI patients with those in health individuals have constantly shown several degrees of cortical “expansion” towards the deafferented brain region and cortical activity in the deprived area with divergent results regarding the appearance of neuropathic pain (Moxon et al., 2014). Similar results are also observed in amputees in which phantom limb pain has been either associated with a stronger representation of the missing body part and maintenance of its structural integrity (Makin et al., 2013) or a remapping of the cortical areas (Flor et al., 1995). It is thought that the degree of cortical reorganization can be attributed to several variables including species, age, behavior activity and therapies regimes after the injury (Moxon et al., 2014; Jutzeler et al., 2019). Since we did not find any differences regarding age, sex or anatomical location of the electrode, we postulate that the population that did not exhibit changes in the cortical activity may result from our acute model of SCI (i.e. from minutes to few hours), which does not exclude the possibility that these animals present cortical changes in sub-acute and/or chronic phases of the injury. In addition, it would be interesting in future to determine whether such small population is directly linked, or not, to sensorimotor pathologies usually observed after SCI. Although our main results are supported by the consistent effects observed in the 79% of animals with SCI that we have studied, we found these results very important as they increase the variability and heterogeneity of effects observed in individuals with SCI, which should be taken into account in the context of cortical reorganization.

## ACKNOWLEGMENTS

We thank Dr. Casto Rivadulla for helpful comments on the manuscript. This work was supported by Spanish Ministry of Economy and Competitiveness and Ministry of Science, Innovation and Universities co-funded by FEDER to J.A. (BFU2016-80665-P and PID2019-105020GB), to A.O. (SAF2016-80647-R), to M.Z (BES2017-082029, predoctoral fellowship FPI-MICINN) and to J.M.R. (RYC2019-026870-I). JMR was also funded by European Union’s Horizon 2020 research and innovation programme under the Marie Sklodowska-Curie grant agreement N° 794926.

## AUTHOR CONTRIBUTIONS

Conceptualization, M.Z, J.M.R., J.A.; Methodology, M.Z., J.M.R., C.M-Q., E.A-C., E.F-L., J.A.; Investigation, M.Z., J.M.R., C.M-Q., E.A-C., E.F-L.; Original Draft, J.M.R., M.Z., J.A.; Writing–Review & Editing, M.Z., C.M-Q., J.M.R., E.A-C., E.F-L., A.O., J.A.; Visualization, M.Z., J.M.R., J.A.; Funding Acquisition, J.M.R. A.O., J.A.

---

**Supplementary Table 1:**
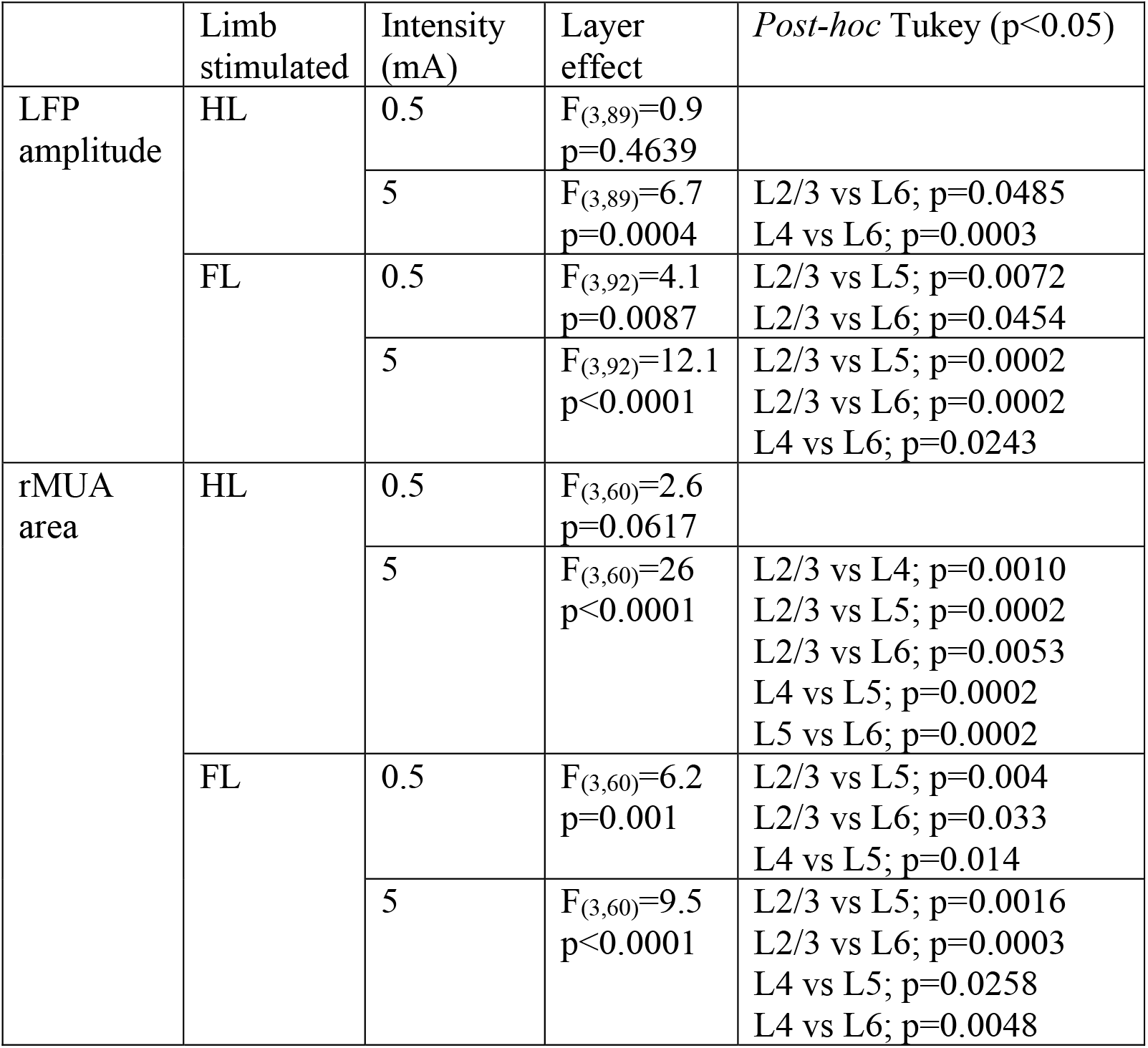
Related to Figure 1, Evoked responses in HLCx by hindlimb and forelimb stimulation. One-way Analysis of Variance (ANOVA) for control responses. Tukey post hoc comparisons are shown for significances p < 0.05.

| | Limb stimulated | Intensity (mA) | Layer effect | *Post-hoc* Tukey ($p < 0.05$) |
| --- | --- | --- | --- | --- |
| LFP amplitude | HL | 0.5 | $F_{(3,89)}=0.9$<br>$p=0.4639$ | |
| | | 5 | $F_{(3,89)}=6.7$<br>$p=0.0004$ | L2/3 vs L6; $p=0.0485$<br>L4 vs L6; $p=0.0003$ |
| | FL | 0.5 | $F_{(3,92)}=4.1$<br>$p=0.0087$ | L2/3 vs L5; $p=0.0072$<br>L2/3 vs L6; $p=0.0454$ |
| | | 5 | $F_{(3,92)}=12.1$<br>$p<0.0001$ | L2/3 vs L5; $p=0.0002$<br>L2/3 vs L6; $p=0.0002$<br>L4 vs L6; $p=0.0243$ |
| rMUA area | HL | 0.5 | $F_{(3,60)}=2.6$<br>$p=0.0617$ | |
| | | 5 | $F_{(3,60)}=26$<br>$p<0.0001$ | L2/3 vs L4; $p=0.0010$<br>L2/3 vs L5; $p=0.0002$<br>L2/3 vs L6; $p=0.0053$<br>L4 vs L5; $p=0.0002$<br>L5 vs L6; $p=0.0002$ |
| | FL | 0.5 | $F_{(3,60)}=6.2$<br>$p=0.001$ | L2/3 vs L5; $p=0.004$<br>L2/3 vs L6; $p=0.033$<br>L4 vs L5; $p=0.014$ |
| | | 5 | $F_{(3,60)}=9.5$<br>$p<0.0001$ | L2/3 vs L5; $p=0.0016$<br>L2/3 vs L6; $p=0.0003$<br>L4 vs L5; $p=0.0258$<br>L4 vs L6; $p=0.0048$ |

**Supplementary Table 2.** Related to Figure 3, Evoked responses in deafferented HLCx by forelimb stimulation. Two-way repeated measures Analysis of Variance (ANOVA) for comparisons between pre- and post-lesion responses across layers when forelimb was stimulated. Post hoc significance is described just for comparisons between conditions in the same layer (lesion effect).

|  | Group | mA | Lesion effect | Layer effect | Lesion* Layer effect | Post-hoc Tukey |
| --- | --- | --- | --- | --- | --- | --- |
| LFP amplitude | All | 0.5 | $F_{(1,92)}=51.8$<br>$p<0.0001$ | $F_{(3,92)}=3.2$<br>$p=0.0256$ | $F_{(3,92)}=0.2$<br>$p=0.9198$ | L2/3 $p=0.0555$<br>L4 $p=0.0059$<br>L5 $p=0.0033$<br>L6 $p=0.0150$ |
| | | 5 | $F_{(1,91)}=50.2$<br>$p<0.0001$ | $F_{(3,91)}=10.3$<br>$p<0.0001$ | $F_{(3,91)}=0.3$<br>$p=0.8074$ | L2/3 $p=0.0227$<br>L4 $p=0.0019$<br>L5 $p=0.0051$<br>L6 $p=0.1130$ |
| rMUA area | All | 5 | $F_{(1,60)}=18.1$<br>$p<0.0001$ | $F_{(3,60)}=12.5$<br>$p<0.0001$ | $F_{(3,60)}=1.2$<br>$p=0.3145$ | L2/3 $p=0.9945$<br>L4 $p=0.6444$<br>L5 $p=0.0378$<br>L6 $p=0.1315$ |
| LFP amplitude | Group 1 (79%) | 0.5 | $F_{(1,72)}=48.5$<br>$p<0.0001$ | $F_{(3,72)}=2.3$<br>$p=0.0846$ | $F_{(3,72)}=0.2$<br>$p=0.9028$ | L2/3 $p=0.0924$<br>L4 $p=0.0136$<br>L5 $p=0.0048$<br>L6 $p=0.0151$ |
| | | 5 | $F_{(1,72)}=125$<br>$p<0.0001$ | $F_{(3,72)}=7$<br>$p=0.0003$ | $F_{(3,72)}=0.5$<br>$p=0.6790$ | L2/3 $p=0.0004$<br>L4 $p=0.0001$<br>L5 $p=0.0001$<br>L6 $p=0.0002$ |
| rMUA area | Group 1 (79%) | 5 | $F_{(1,47)}=25.5$<br>$p<0.0001$ | $F_{(3,47)}=9.5$<br>$p<0.0001$ | $F_{(3,47)}=1$<br>$p=0.4020$ | L2/3 $p=0.9101$<br>L4 $p=0.3758$<br>L5 $p=0.0148$<br>L6 $p=0.0643$ |
| LFP amplitude | Group 2 (21%) | 0.5 | $F_{(1,16)}=4.5$<br>$p=0.050$ | $F_{(3,16)}=0.8$<br>$p=0.5111$ | $F_{(3,16)}=0.1$<br>$p=0.9493$ | |
| | | 5 | $F_{(1,15)}=15.2$<br>$p=0.0014$ | $F_{(3,15)}=4.2$<br>$p=0.0250$ | $F_{(3,15)}=1.8$<br>$p=0.1826$ | L2/3 $p=0.9998$<br>L4 $p=0.9085$<br>L5 $p=0.1163$<br>L6 $p=0.1218$ |
| rMUA area | Group 2 (21%) | 5 | $F_{(1,8)}=0.5$<br>$p=0.5111$ | $F_{(3,8)}=4.1$<br>$p=0.0492$ | $F_{(3,8)}=0.1$<br>$p=0.9725$ | |

**Supplementary Table 3:**
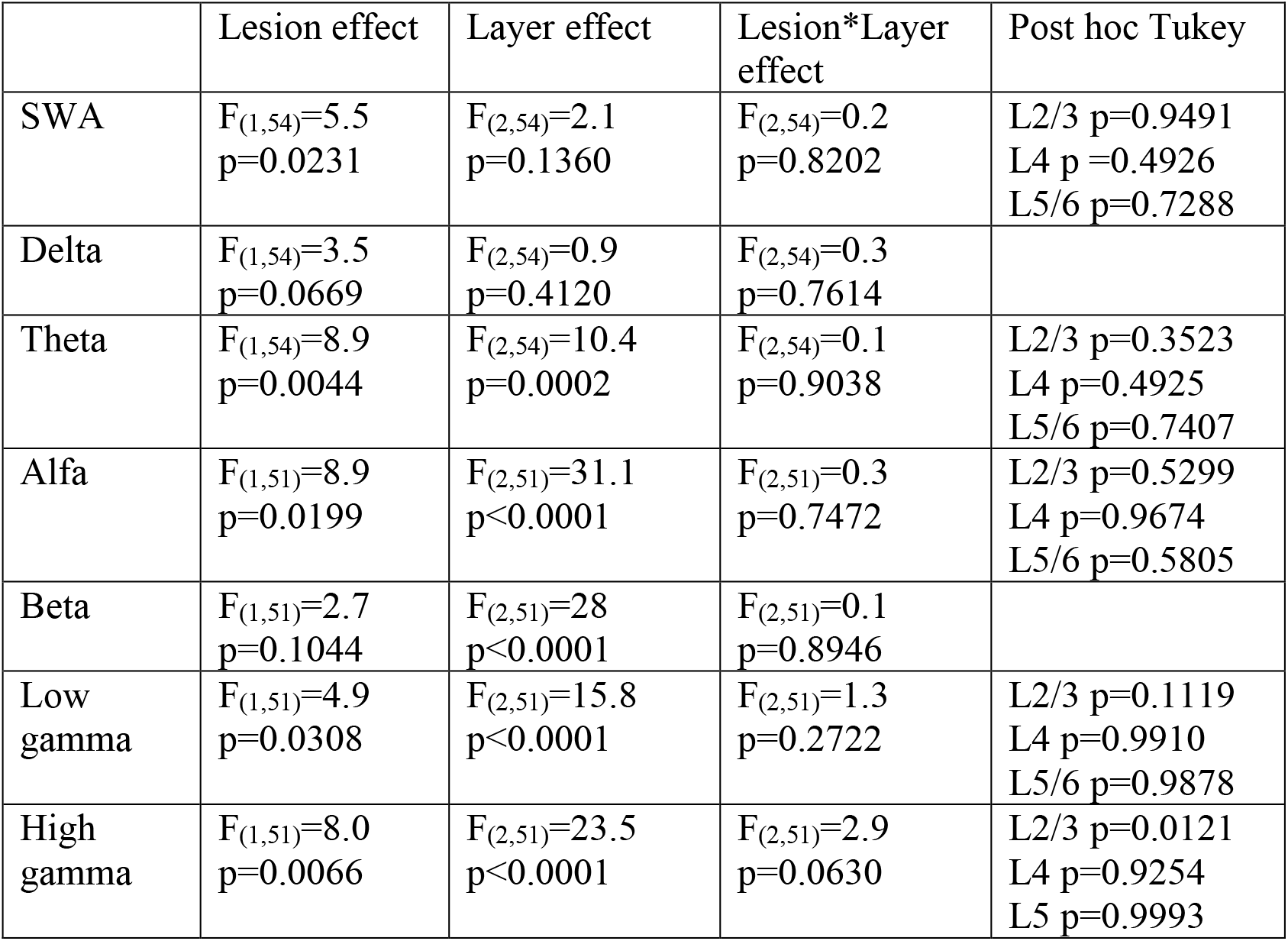
Related to Figure 8, Frequency content across layers of HLCx before and immediately after a SCI. Two-way Analysis of Variance (ANOVA) of the relative power spectrum of each band frequency in the spontaneous activity fast transform fourier analysis. Post hoc significance is described just for comparisons between conditions in the same layer (lesion effect). Analysis from Group 1 animals.

|  | Lesion effect | Layer effect | Lesion*Layer effect | Post hoc Tukey |
| --- | --- | --- | --- | --- |
| SWA | $F_{(1,54)}=5.5$<br>$p=0.0231$ | $F_{(2,54)}=2.1$<br>$p=0.1360$ | $F_{(2,54)}=0.2$<br>$p=0.8202$ | L2/3 $p=0.9491$<br>L4 $p=0.4926$<br>L5/6 $p=0.7288$ |
| Delta | $F_{(1,54)}=3.5$<br>$p=0.0669$ | $F_{(2,54)}=0.9$<br>$p=0.4120$ | $F_{(2,54)}=0.3$<br>$p=0.7614$ | |
| Theta | $F_{(1,54)}=8.9$<br>$p=0.0044$ | $F_{(2,54)}=10.4$<br>$p=0.0002$ | $F_{(2,54)}=0.1$<br>$p=0.9038$ | L2/3 $p=0.3523$<br>L4 $p=0.4925$<br>L5/6 $p=0.7407$ |
| Alfa | $F_{(1,51)}=8.9$<br>$p=0.0199$ | $F_{(2,51)}=31.1$<br>$p<0.0001$ | $F_{(2,51)}=0.3$<br>$p=0.7472$ | L2/3 $p=0.5299$<br>L4 $p=0.9674$<br>L5/6 $p=0.5805$ |
| Beta | $F_{(1,51)}=2.7$<br>$p=0.1044$ | $F_{(2,51)}=28$<br>$p<0.0001$ | $F_{(2,51)}=0.1$<br>$p=0.8946$ | |
| Low gamma | $F_{(1,51)}=4.9$<br>$p=0.0308$ | $F_{(2,51)}=15.8$<br>$p<0.0001$ | $F_{(2,51)}=1.3$<br>$p=0.2722$ | L2/3 $p=0.1119$<br>L4 $p=0.9910$<br>L5/6 $p=0.9878$ |
| High gamma | $F_{(1,51)}=8.0$<br>$p=0.0066$ | $F_{(2,51)}=23.5$<br>$p<0.0001$ | $F_{(2,51)}=2.9$<br>$p=0.0630$ | L2/3 $p=0.0121$<br>L4 $p=0.9254$<br>L5 $p=0.9993$ |

